# HSV-1 miRNAs are posttranscriptionaly edited in latently infected human ganglia

**DOI:** 10.1101/2023.05.26.542484

**Authors:** Andreja Zubković, Cristina Gomes, Adwait Parchure, Mia Cesarec, Antun Ferenčić, Filip Rokić, Hrvoje Jakovac, Abigail L. Whitford, Sara A. Dochnal, Anna R. Cliffe, Dražen Cuculić, Angela Gallo, Oliver Vugrek, Michael Hackenberg, Igor Jurak

## Abstract

Viruses use miRNAs to enable efficient replication, control host defense mechanisms, and regulate latent infection. Herpes simplex virus 1 (HSV-1) expresses multiple miRNAs, whose functions are largely unknown. The evolutionary conservation of many HSV-1 miRNAs in the closely related HSV-2 suggests their functional importance. miRNAs, similar to other transcripts, can undergo various posttranscriptional modifications that may affect their biogenesis, stability and targeting. To investigate whether editing occurs in HSV-1 miRNAs, we sequenced samples from latently infected human ganglia. We show that one of the six HSV-1 miRNAs (miR-H2 to -H8) that define HSV-1 latency, miR-H2, exhibits A-to-I hyperediting within the miRNA seed sequence. We observed the same specific miR-H2 hyperediting phenomenon in miRNAs isolated from the ganglia of latently infected mice and, to a lesser extent, during productive infection in cultured cells. Curiously, we found no evidence of editing of the encoded HSV-2 homolog in latently infected mice or in cultured cells. The efficient loading of the edited miRNAs onto the RISC complex, indicates their ability to function as miRNAs. To investigate the potential of the edited miRNA to alter mRNA targeting, we predicted the host and viral targets for the modified miRNAs. Nucleotide substitution in the seed region significantly increased the number of potential host and viral targets. Most notably, ICP4, an essential viral protein, was predicted to be an additional target. Using transfection assays, we demonstrated that edited miRNAs have the potential to regulate ICP4 in addition to the previously identified target ICP0. Our study identifies a specific hyperedited HSV-1 mRNA, miR-H2, and highlights how the virus can use a single miRNA to target multiple transcripts during persistent, latent infection.

**Importance:** Herpes simplex virus 1 is an important human pathogen and intensively studied for many decades. Nevertheless, the molecular mechanisms regulating its establishment, maintenance, and reactivation from latency are poorly understood. Here, we show that HSV- 1 encoded miR-H2 is post-transcriptionally edited in latently infected human tissues. Hyperediting of viral miRNAs increases the targeting potential of these miRNAs and may play an important role in regulating latency. We show that the edited miR-H2 (miR-H2-e) can target ICP4, an essential viral protein. Interestingly, we found no evidence of hyperdating of its homolog, miR-H2, which is expressed by the closely related virus HSV-2. The discovery of posttranslational modifications of viral miRNA in the latency phase suggests that these processes may also be important for other non-coding viral RNA in the latency phase, including the intron LAT, which in turn may be crucial for understanding the biology of this virus.

## Introduction

miRNAs are small non-coding RNAs expressed by plants, animals and viruses. miRNAs regulate gene expression mainly by binding to partially complementary transcripts and affecting their stability and translation. The specificity of miRNA binding to its target is largely determined by a short sequence at the 5’ end of the miRNA molecule (nucleotides 2- 8) called the seed (reviewed in (1–3)). miRNAs regulate most protein-coding genes and are essential for normal development and growth. On the other hand, deregulated miRNAs are associated with a number of diseases including cancer (4).

Viruses, particularly large DNA viruses, have been found to encode or use host miRNAs to regulate their infection (5–7). Herpes simplex virus 1 (HSV-1) encodes miRNAs from 20 loci (miR-H1 – H18, -H26-H29), some of which are conserved in their genomic position and some of which are partially conserved in their sequence in the closely related herpes simplex virus 2 (HSV-2) (8–13). The genomic miRNA loci are scattered throughout the HSV-1 genome, but the most abundantly expressed miRNAs (miR-H1 to -H8) are located in close proximity or within the region encoding the latency associated transcripts (LATs)(7).

LATs are group of long non-coding RNAs abundantly expressed during latent infection, whose function is quite enigmatic. Nonetheless, several functions have been ascribed to the LATs, including: inhibition of apoptosis, repression of productive gene expression and viral chromatin organization (14, 15). The expression of miRNAs encoded within the LATs is strongly dependent on the activity of the LAT promoter (miR-H2-H8), although other promoters may also contribute to their expression (16–18). These miRNAs follow the kinetics of LATs in establishment, maintenance, and reactivation from latency (16, 19, 20). It has been shown that miRNAs encoded upstream and in close proximity to the LAT promoter, miR-H1 and miR-H6, can negatively regulate the LAT transcript and LAT associated miRNAs during productive infection in cell culture (21). Many of the LATs associated miRNAs are encoded antisense to key viral genes and are fully complementary to their transcripts, suggesting regulation of these genes (7). For example, miR-H2, -H7 and -H8 are encoded antisense to ICP0, a viral ubiquitin ligase that directs many proteins for degradation and has a function in inhibiting host defense mechanisms and promoting viral transcriptional activation (22, 23). miR-H3 and -H4 are encoded antisense to ICP34.5, also an inhibitor of intrinsic innate immunity (24). Viral mutants deficient for the expression of individual miRNAs usually exhibit little or no replication defects in cultured cells and retain wild-type or mild latency reactivation phenotype in animal models (21, 25–27). Remarkably, in cultured cells miR-H28 and -H29 have been shown to be exported by exosomes and that can limit virus transmission in recipient cells by inducing IFN production (12, 28). In addition to expressing its own miRNAs, HSV-1 deregulates the host miRNAome, and host miRNAs have been shown to contribute to efficient viral infection (29). For example, the neuronal miRNA miR-138 regulates the expression of ICP0 and multiple host targets to promote latent infection (30, 31). Nonetheless, exploring the exact roles of miRNAs in HSV-1 latency presents many challenges and has yet to be discovered.

Adenosine Deaminase Acting on RNA (ADAR) is a family of editing enzymes that catalyze adenosine-to-inosine (A-to-I) deamination in dsRNA (32). Inosine is read as guanosine by the translational machinery and viral RNA dependent RNA polymerases, leading to recoding events (modified codons) and changes in viral genome (A-to-G transition). In addition, adenosine deamination can affect translational efficiency, change splicing, RNA structure and stability, and miRNA targeting (32). In humans, editing occurs at millions of sites and mostly in non-coding parts of the transcriptome (introns and untranslated regions UTRs) typically targeting dsRNA formed by pairing of two inverted copies of repetitive elements (33). The vast majority of A-to-I editing activity occurs within ∼300 base-pair long Alu repeats represents in more than a million copies. Vertebrates have three ADAR genes, ADAR1, ADAR2 and ADAR3. ADAR1 and ADAR2 are enzymatically active, whereas ADAR3 has been shown to have a regulatory role (32). ADAR1 is expressed in two isoforms, a shorter isoform (ADAR1-p110) expressed from a constitutively active promoter, and ADAR-p150 which is interferon inducible. The ADAR1 mediated editing has essential role in preventing activation of innate immune system and interferon overproduction by endogenous dsRNA (32).

Roles of ADAR proteins in virus replication, including herpesviruses, are not well understood and there is evidence for antiviral and pro-viral properties depending on virus and host (34, 35). Editing of viral transcripts during productive or latent infection has been documented for some herpesviruses potentially affecting important biological processes (34, 36, 37). For example, editing of one of the most abundant transcripts found in KSHV latency, K12 (kaposin A) open reading frame, eliminates its transforming activity (38). More recently, Rajendran et al. found a number of additional edited transcripts, including KSHV encoded miR-K12-4-3p (39). Editing of this miRNA impacts miRNA biogenies and its target specificity. Moreover, ADAR1 function was found required for efficient KSHV lytic reactivation by modulating innate immunity signaling (40). Similarly, EBV encoded miRNAs and other latent transcripts have been found edited (41–43). Interestingly, editing of ebv-miR-BART6 and - BART3 reduces their Dicer targeting and affects viral latency. On the other hand, HCMV infection triggers increased ADAR1 levels and enhanced editing of host miR-376a, which in turn acquires the ability to downregulate the immunomodulatory molecule HLA-E and facilitate immune elimination of HCMV infected cells (44).

In this study, we report a comprehensive analysis of HSV-1 miRNAs in latently infected human ganglia. We found evidence of specific A-to-I hyperediting of an HSV-1 miRNA, miR-H2. miR-H2 is hyperedited at nucleotide position N5 within the seed region, indicating a change in its specificity and targeting. We provided evidence that the edited miR-H2 has additional targets and that it can regulate ICP4, in addition to the previously determined target, ICP0.

## Results

### miRNA sequencing

To analyze the expression of HSV-1 miRNAs in latently infected human neurons we sequenced small RNAs extracted from sections of ten latently infected human trigeminal ganglia (TGs) (Supp. Table 1). Briefly, we extracted RNA and DNA from all samples and confirmed the presence of the HSV-1 genome prior to sequencing (Supp. Fig. 1). Small RNA libraries were generated using the NEBNext Multiplex Small RNA Library PreSet for Illumina and sequenced on the NextSeq 500 (Illumina). The obtained sequencing datasets were processed with sRNAtoobox (45), an integrated collection of small RNA research tools, applying stringent parameters with exclusion of all bases with Phred score <30.

We obtained between 2.6x10^7^ and 1.5x10^7^ high quality sequencing reads per sample, of which between 2,6 x10^6^ and 9,5 x10^6^ represented miRNAs (Table 1). As expected, the vast majority of reads were human miRNAs (>99,8%) and only a small number of HSV-1 miRNA were obtained (2 - 6884 reads/sample) (Table 1). HSV-1 miRNAs detected in each individual sample did not correlate with the number of human miRNAs detected and ranged from 0.00003% to 0.17% (Table 1). These results could indicate differences in viral load, differences in processed sections of individual TG, and others.

**Table 1.**
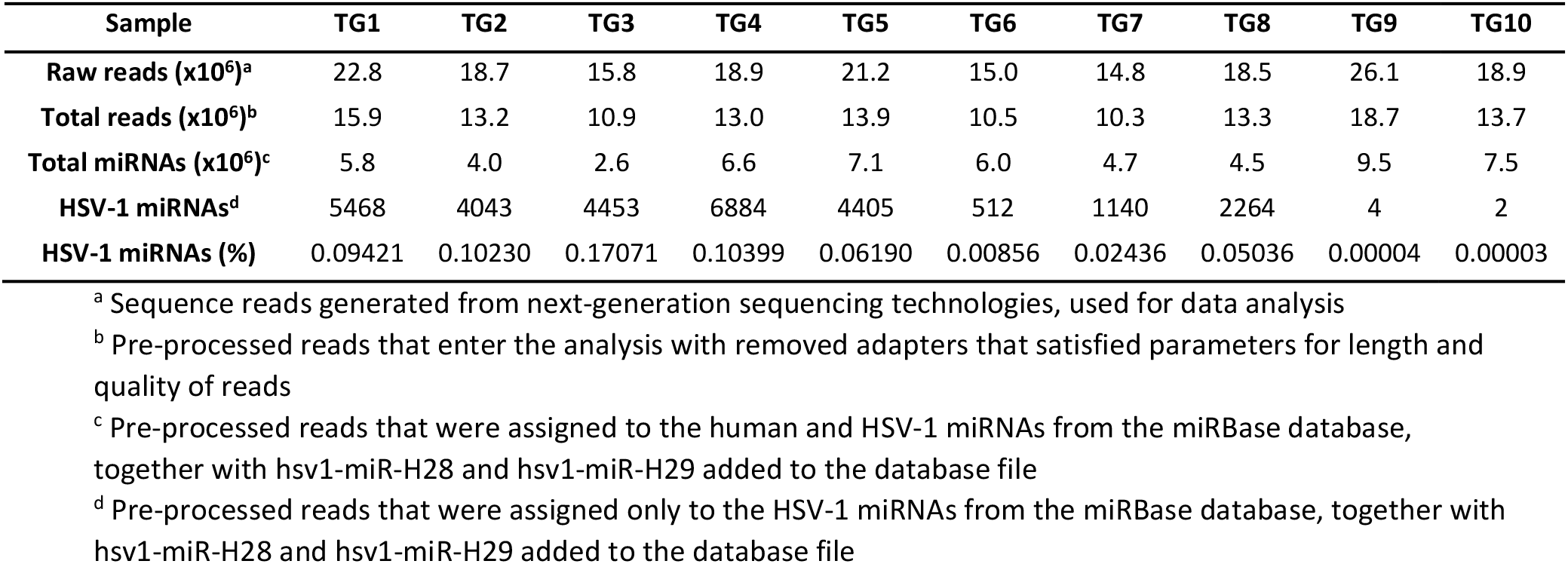
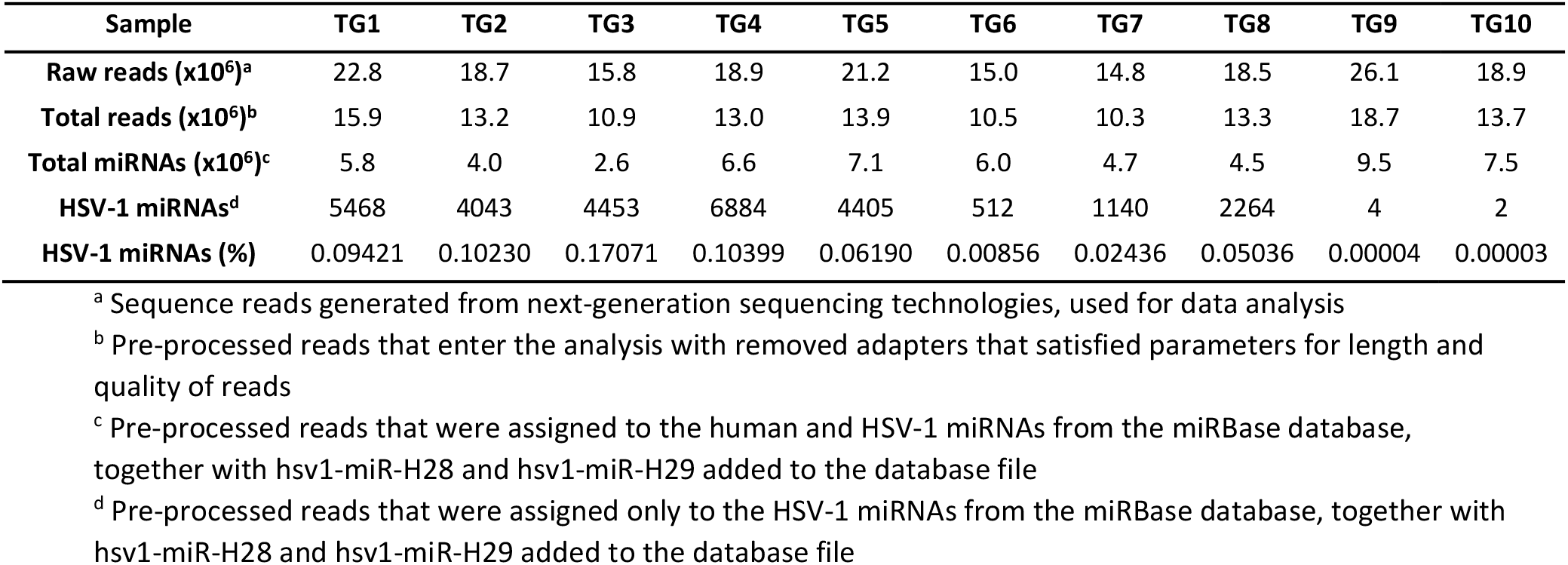
Number of sequencing reads representing human and HSV-1 miRNAs in human trigeminal ganglia.

**Table 2.**
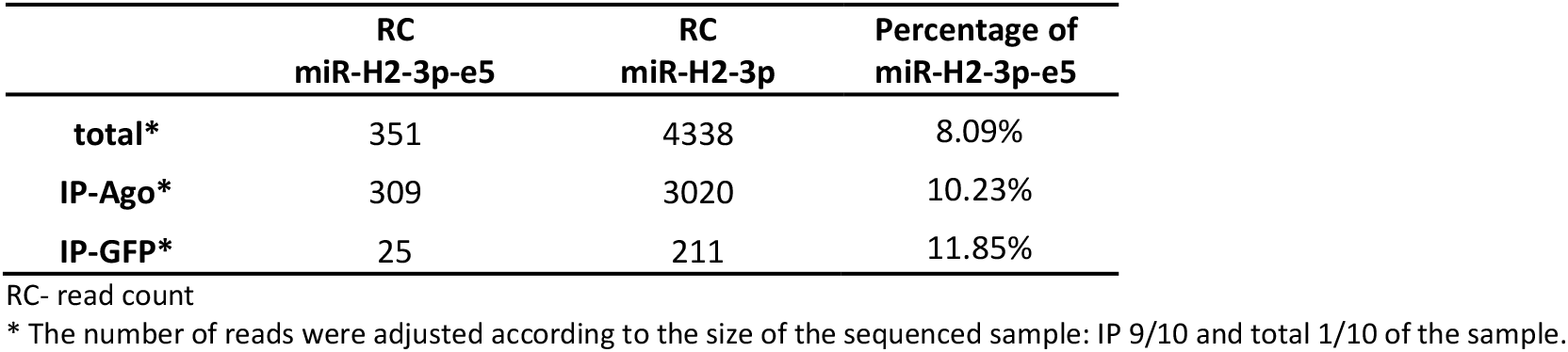
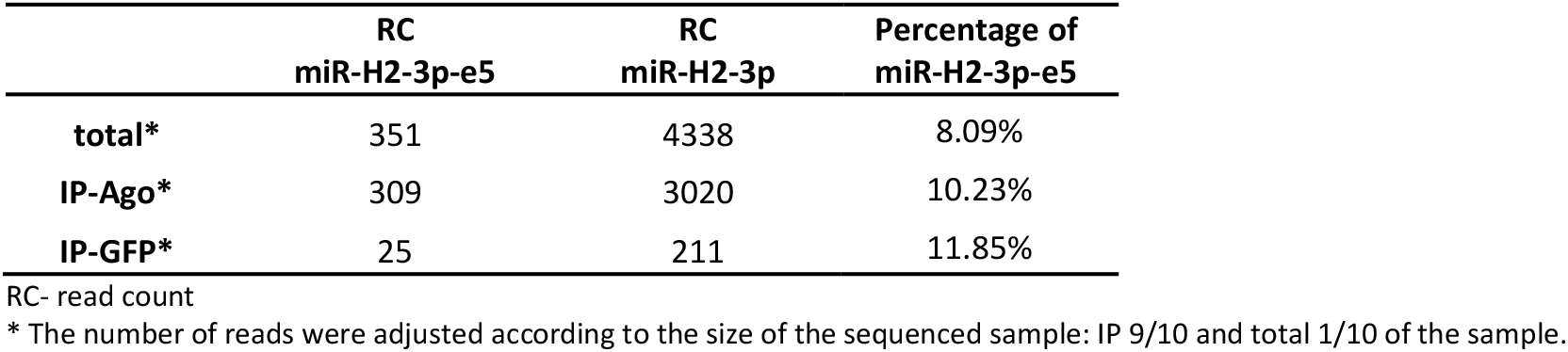
Edited miR-H2 is associated with Ago2.

### The pattern of HSV-1 miRNA expression in human trigeminal ganglia

The expression of HSV-1 miRNAs in latently infected human ganglia has been comprehensively studied in only a few cases (10, 46), and in contrast to studies in cultured cells and animal models, only a small subset of miRNAs has been detected (miR-H2 - miR- H8). Our study shows the expression of these miRNAs in all samples (Figure 1.) with a remarkably similar pattern. In two samples we detected a low number of HSV-1 miRNAs, which could indicate a lower viral load or a variation within each tissue section. Based on the number of reads, the most abundant miRNA was miR-H4 followed by -H3, -H2, -H7, -H6, -H8 and -H5, with a range of 348 reads per million (RMP) for the most abundant and <1 RPM for the least abundant miRNA. The number of reads does not necessarily correspond to absolute quantification, as these numbers may depend heavily on the sequence of the miRNA and the sequencing protocol (47, 48). Nonetheless, these LAT associated miRNAs define the pattern of HSV-1 miRNA in latently infected human trigeminal ganglia. In addition to these miRNAs, we detected miR-H1 and miR-H17 in TG4 and TG7, respectively (Figure 1.). miR-H1 was detected with only 3 reads (0,23 RPM) in sample TG4 but the reads fully matched the miR-H1 sequence deposited in miRbase, indicating a confident result. Since miR-H1 is abundantly expressed during productive infection (9), this could suggest a possible reactivation from latency in one of the samples. On the other hand, miR-H17 was detected with only a single read and the sequence was shorter (19nt) and had a mutation compared to miRNA in miRBase, which may indicate false detection. It is important to note that the abundantly expressed HSV-1 miRNAs (miR-H2, -H3, -H4, -H7, except miR-H6) were readily detected with both strands of the duplex (5p and 3p) in ratios of 1:4 to 1:100, which is well above the average host miRNA strand bias (>1:1000). These results suggest, at least for some HSV-1 miRNA, that both arms of the duplex may regulate specific targets. Indeed, this hypothesis is supported by the analysis of RISC-associated HSV-1 miRNAs in which both strands of the duplex were associated with the RISC complex ((27), and not shown).

**Figure 1.**
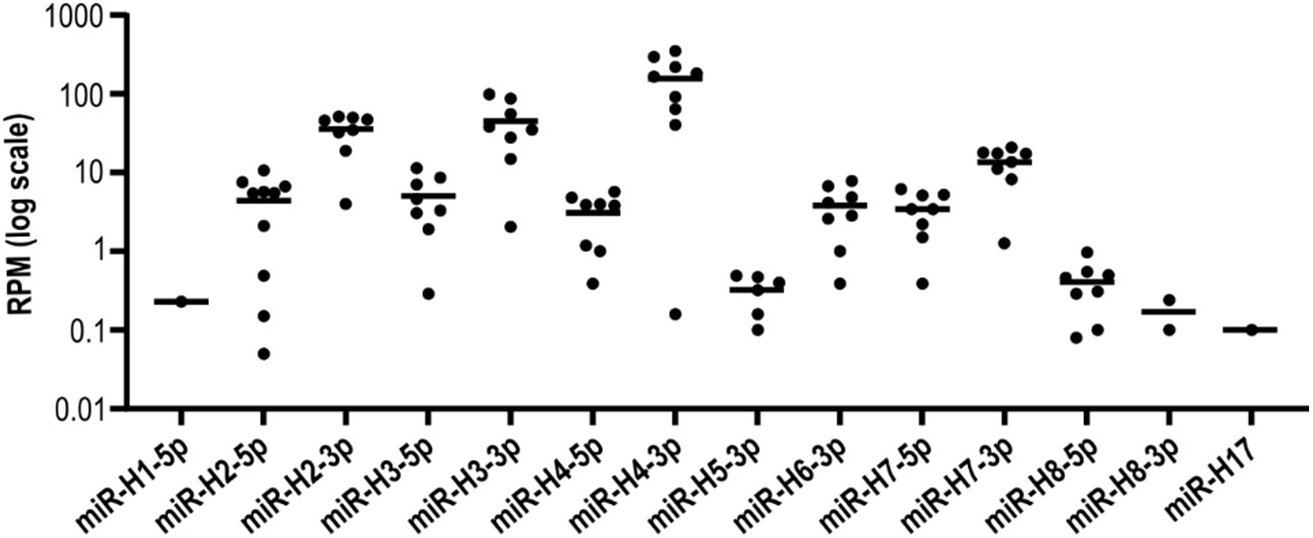
The HSV-1 miRNA expression pattern in human trigeminal ganglia. RNA was extracted from latently infected trigeminal ganglia and small RNA libraries were generated using the NEBNext Multiplex Small RNA Library Pre Set for Illumina and sequenced on the NextSeq 500 (Illumina). The obtained sequencing data sets were processed using sRNAtoolbox. The dots represent the number of reads per million (RPM) for the miRNAs detected in individual ganglia.

### miR-H2 is hyperedited in latently infected human neurons

In our previous study, we identified particular sequence variants of HSV-1 miRNAs that could result from active posttranscriptional modifications by ADAR enzymes (46). To investigate this further, we analyzed the sequence variants of HSV-1 miRNAs in our dataset. In total, we obtained approximately 29 000 reads representing HSV-1 miRNAs (ranging from 2 to 6884/sample) under our threshold settings (i.e. maximum 2 mismatches, allowing 5’ and 3’ fluctuations). Detailed analysis of the reads showed that 13.1% of all miRNAs had 1 or two nucleotide substitutions. This is comparable to the substitutions detected in the host miRNA dataset (i.e. 11,5%). Remarkably, among all substitutions within viral miRNAs, we observed a very high frequency of A-to-G substitutions, which accounted for more than 50% of all nucleotide substitutions (Figure 2A). Importantly, the frequency of nucleotide substitutions was comparable between host and viral miRNAs, and most types of substitutions were relatively rare. However, A-to-G substitutions were the most common substitution in both host and virus, and surprisingly, they occurred twice as frequently in virus as in host (Figure 2B). It is possible that some of detected substitutions are sequencing errors, but the observed high frequency of A-to-G substitutions indicate an active enzymatic process. Next, we carefully analyzed the A-to-G substitutions for each miRNA, and found that the frequency of A-to-G substitutions for most viral miRNAs was very low, similar to the baseline level (<1%) (Figure 2C). On the other hand, miR-H2-3p was in each sample represented with 35% to 48% reads having A-to-G substitutions and accounted for nearly 80% of all A-to-G substitutions in HSV-1 miRNAs (Figure 2C and D). We observed the vast majority of A-to-G substitutions at the nucleotide positions N5 (miR-H2-3p-e5) (Figure 2D). In addition, we detected lower level of editing at the N9 nucleotide position (<10%). The pattern of N5 and N9 nucleotide substitution was conserved in all latently infected TG with detectable miR-H2 (Figure 2E). Other types of substitutions were at negligible levels (Figure 2D).

**Figure 2.**
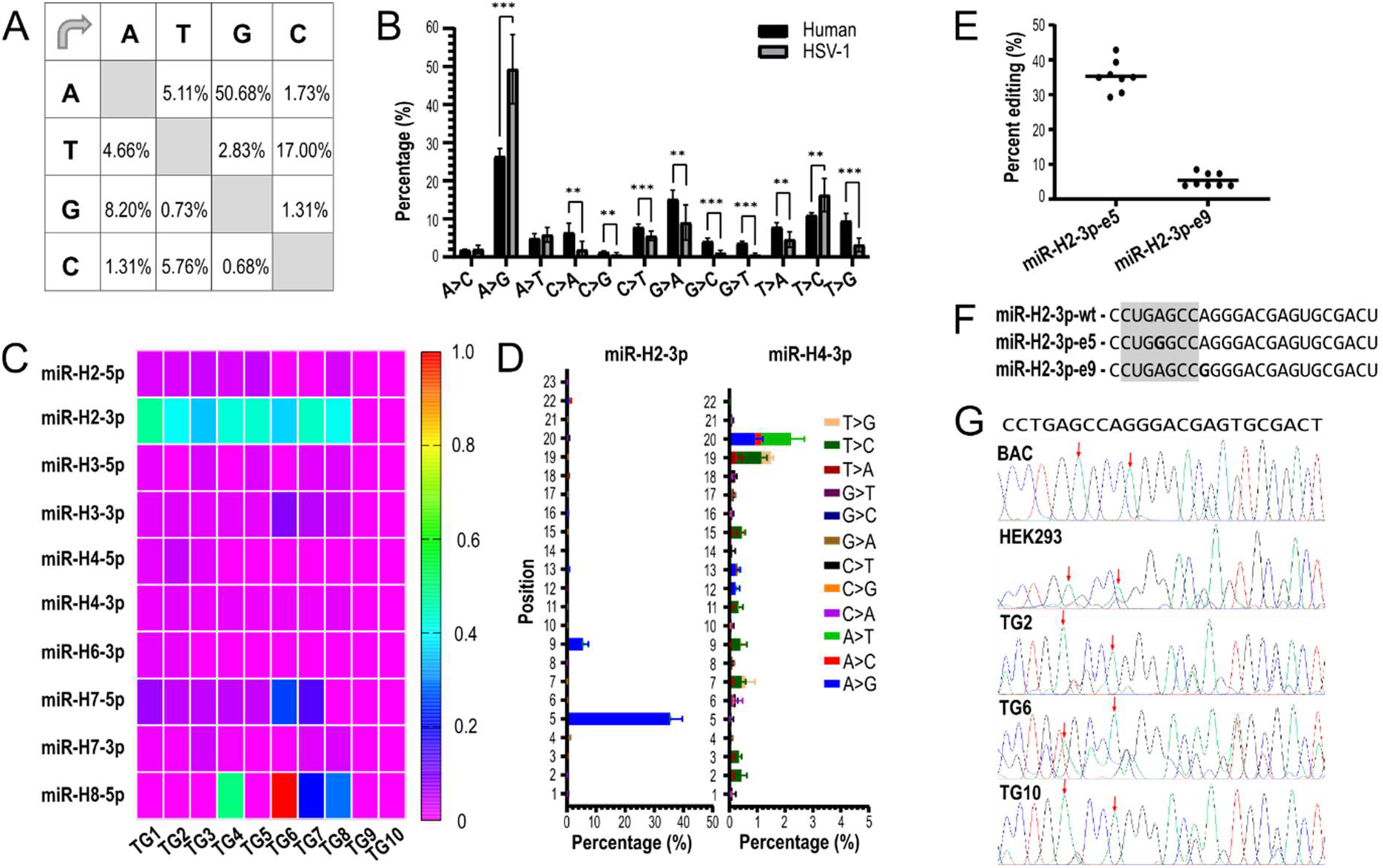
miR-H2 is hyperedited in latently infected human neurons. A) Nucleotide substitution matrix of all HSV-1 miRNAs detected in 8 hTGs (n=29 175 miRNA reads). B) Percentage of substitutions of host and viral miRNAs. C) Each panel represents the frequency of A-to-G substitutions for the miRNAs detected in each sample. D) Percentage of the nucleotide substitutions per nucleotide in miRNA-H2- 3p and miR-H4-3p marked in different colors. E) Percentage of edited miR-H2-3p reads in human TGs. Each dot represents the percentage of indicated miRNA in individual TG sample. miR-H2-3p-e5 and miR-H2-3p-e9 indicate sequence editing at the nucleotide position N5 and N9, respectively (number of all miR-H2-3p reads =1 491). F) Sequences of canonical hsv1-miR-H2-3p and edited isoforms. Edited A-to-G nucleotides are shown in bold at indicated positions N5 and N9. The seed sequence of miR-H2- 3p is highlighted in gray. G) The pre-miR-H2 locus was amplified using 50ng of DNA extracted from three latently infected human ganglia (TG2, TG6 and TG10), the cloned HSV-1 genome strain KOS (BAC) and HEK293 cells productively infected with HSV-1 strain KOS (HEK293) using the primers FP: 5’- GCCCGGCCCCCCCGAG-3’, RP: 5’-ACACGGCGCGCGUCCG-3’; and directly sequenced with FP primer. The red arrows point to the nucleotides at positions N5 and N9 of miR-H2-3p.

This result was quite surprising, as HSV-1 miRNAs are considered highly conserved. Indeed, comparison of the pre-mR H2 sequence of laboratory and clinical strains showed strict conservation (Suppl. Figure 2). Moreover, the reproducible pattern of the miR-H2 editing in all analyzed samples strongly indicates a specific modification. However, to exclude the possibility that the sequence variants were due to heterogeneous infection, i.e. different virus strains carrying nucleotide variations in the miR-H2 gene, we sequenced PCR amplicons generated on the DNA template of three TGs analyzed (TG2, TG6, TG10). We also sequenced amplicons obtained from the cloned genome of the HSV-1 strain KOS and DNA extracted from cells infected with HSV-1 in culture, which served as a monoclonal control. In the case of heterogeneous infection, one would expect overlapping sequencing signals for nucleotides A and G. In contrast, we found no evidence of variation at the indicated editing sites in any of the samples (Fig. 2 E). Overall, our results suggest that miR-H2 variants arise by posttranscriptional sequence-specific editing in latently infected neurons, which is similar to the signature of A-to-I deamination by ADAR proteins. Curiously, we also observed other nucleotide substitutions in viral miRNAs, such as T-to-C (17% overall) or C-to-T (5.7%)(Figure 2 A and D), which were distributed across all detected viral miRNAs with low frequency for individual miRNAs (not shown) and were therefore not analyzed further. Figure 4 D. shows the substitutions per nucleotide in miR-H4, an example of a highly expressed viral miRNA that, unlike miR-H2, shows no signs of hyperediting.

ADAR1 and ADAR2, the two enzymatically active members of the ADAR protein family, are known to be ubiquitously expressed in various tissues, including the brain. To confirm that these genes are expressed in the tissue where we detected potential post-transcriptional editing, we analyzed RNA and proteins extracted from human TG tissue. Using RT-PCR, we confirmed the expression of ADAR1 and ADAR2 in human TGs (Figure 3). On the other hand, we detected higher levels of ADAR1 p110 protein compared with the interferon-inducible p150 form, which could be detected only with an isoform-specific antibody (Figure 3). The expression of ADAR2 was below our detection limit, consistent with relatively low levels of the ADAR transcript. In addition, we detected nuclear and moderate cytoplasmic staining of neuronal cells with immunohistochemistry and two different ADAR1 antibodies. ADAR1 expression was also observed in small surrounding cells that may correspond morphologically to satellite glial cells (two middle panels). ADAR2, on the other hand, was observed almost exclusively in nuclei and was also present in surrounding satellite cells (lower panel). Immunohistochemical signals were not detected in stromal connective tissue. Taken together, these results suggest that ADAR proteins can indeed process miR-H2 in latently infected human TG.

**Figure 3.**
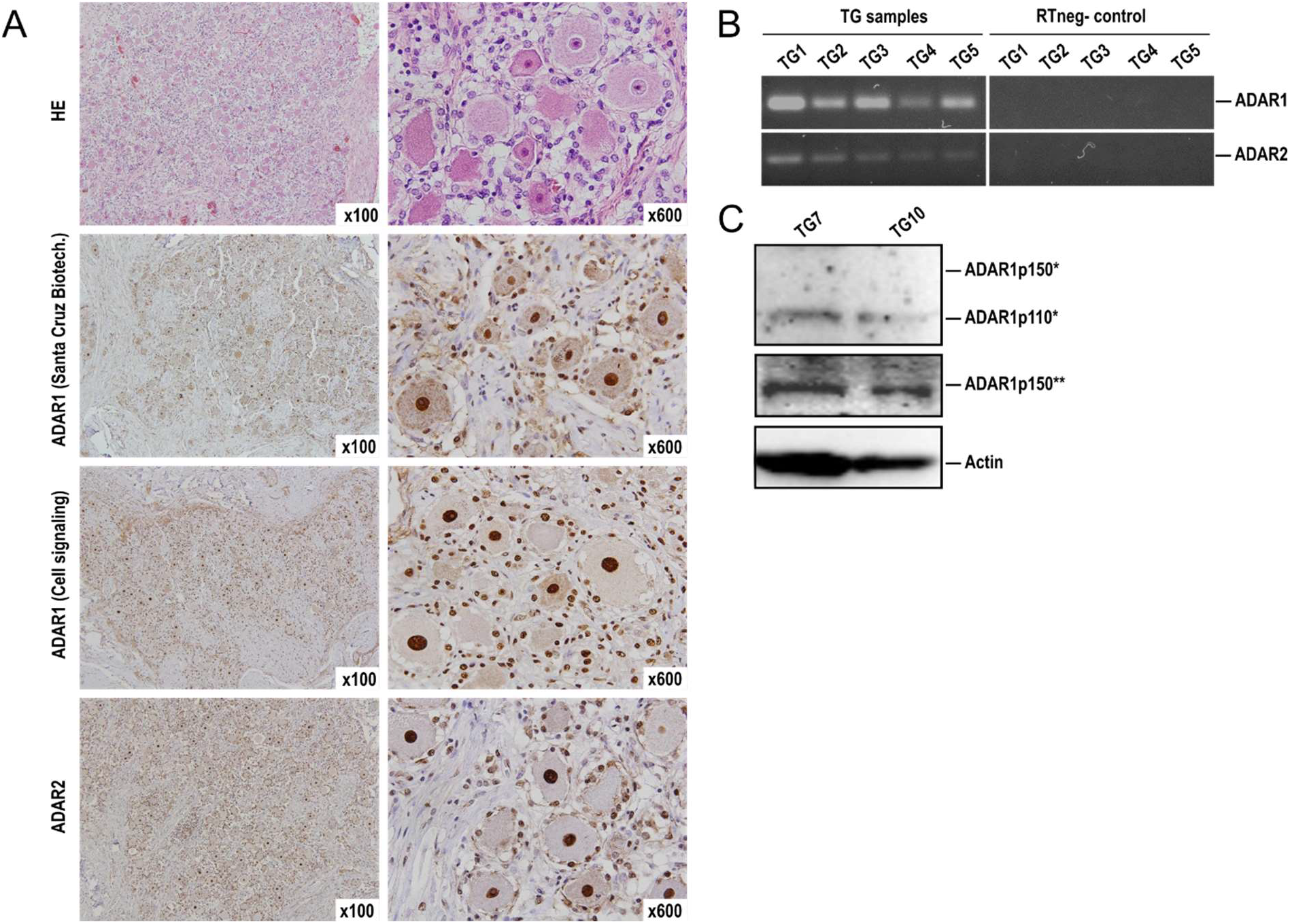
Expression of ADAR1 and ADAR2 in human trigeminal ganglia. A) Immunohistochemical analysis of ADAR1 and ADAR2 expression in human TGs. 4 µm sections of paraffine-embedded TG tissue were stained with rabbit monoclonal α-ADAR1 IgG (Cell Signaling Technology), mouse monoclonal α-ADAR1 IgG (Santa Cruz) and mouse monoclonal α-ADAR2 IgG (Santa Cruz;) and visualized using peroxidase-labeled polymer linked to goat α-rabbit and α-mouse immunoglobulins and 3,3′-Diaminobenzidine (DAB). Slides were counterstained with hematoxylin and analyzed by an Olympus BX51 microscope equipped with a DP50 camera and Cell^F software (Olympus, Japan). The magnification is indicated in each panel (x100 or x600). HE, ADAR1, and ADAR2 staining is shown in neurons; B) RNA was extracted from human trigeminal ganglia (indicated TG samples TG1-TG5) and ADAR1 and ADAR2 transcripts were amplified using RT-PCR and specific primers. PCR products were visualized using by agarose gel electrophoresis. RTneg- control represents the samples processed without the reverse transcriptase.; C) Proteins were isolated from sections of two randomly selected human trigeminal ganglia (TG7 and TG9) with RIPA buffer and analyzed by Western blot. ADAR1 was detected with the α-ADAR1 antibody (Santa Cruz)(upper panel) or the ADAR1 p150 antibody (Cell Signaling) (middle panel).

### Low levels of HSV-1 miRNA editing during productive infection in culture

We demonstrated that sequence-specific hyperediting of miR-H2 occurs during long-lasting latency in humans, which might have important implications for understanding the molecular mechanisms regulating latency. We next asked whether editing of miR-H2 could also be observed during productive infection in cultured cells, which would strongly suggest sequence-specific deamination. Importantly, under such experimental conditions, the infection can be considered monoclonal with respect to sequence variations. To address these questions, we reanalyzed our previous sequencing of smRNAs from productively infected HFF with the HSV-1 strain KOS at an MOI of 5 (ref) at 8 and 18 hour post infection (h.p.i.). As expected, we detected almost all previously reported HSV-1 miRNAs (except miR- H3 -H18 and -H27), which differed significantly in abundance (Figure 4A). HSV-1 miRNAs were represented with 243 714 reads and 8.71 % had one or two nucleotide substitutions, consistent with the basal frequency of substitutions observed in human latency samples. We detected high levels of A-to-G substitutions (12,4%), which was significantly higher than most of other substitutions, but still the phenomenon of hyperediting was less pronounced than in latently infected human ganglia (Figure 4B). In contrast to latency in humans, miR-H2 editing was not highly overrepresented (14 % of all A-to-G editing) and it was slightly above the frequency of A-to-G substitutions in most detected miRNAs. However, we observed two important correlations with the latent samples. First, miR-H2 editing showed a slight but reproducible increase from early (8 h) to late (18 h) times postinfection, i.e. from 2,7% to 5,9% (Figure 4 D). And second, miR-H2 editing occurred at the same nucleotides (N5 and N9) and in similar ratios to latency (Figure 4C). In addition, we observed higher frequency of A-to-G substitutions for miR-H11, -H12 and -H13, miRNAs that arise from transcripts spanning oriS and oriL regions (Fig. 4C). Although we cannot determine whether these substitutions represent true editing events, due to the very small number of reads, it is possible that palindromic sequences of oriS and oriL could be genuine substrates for ADAR binding and editing.

**Figure 4.**
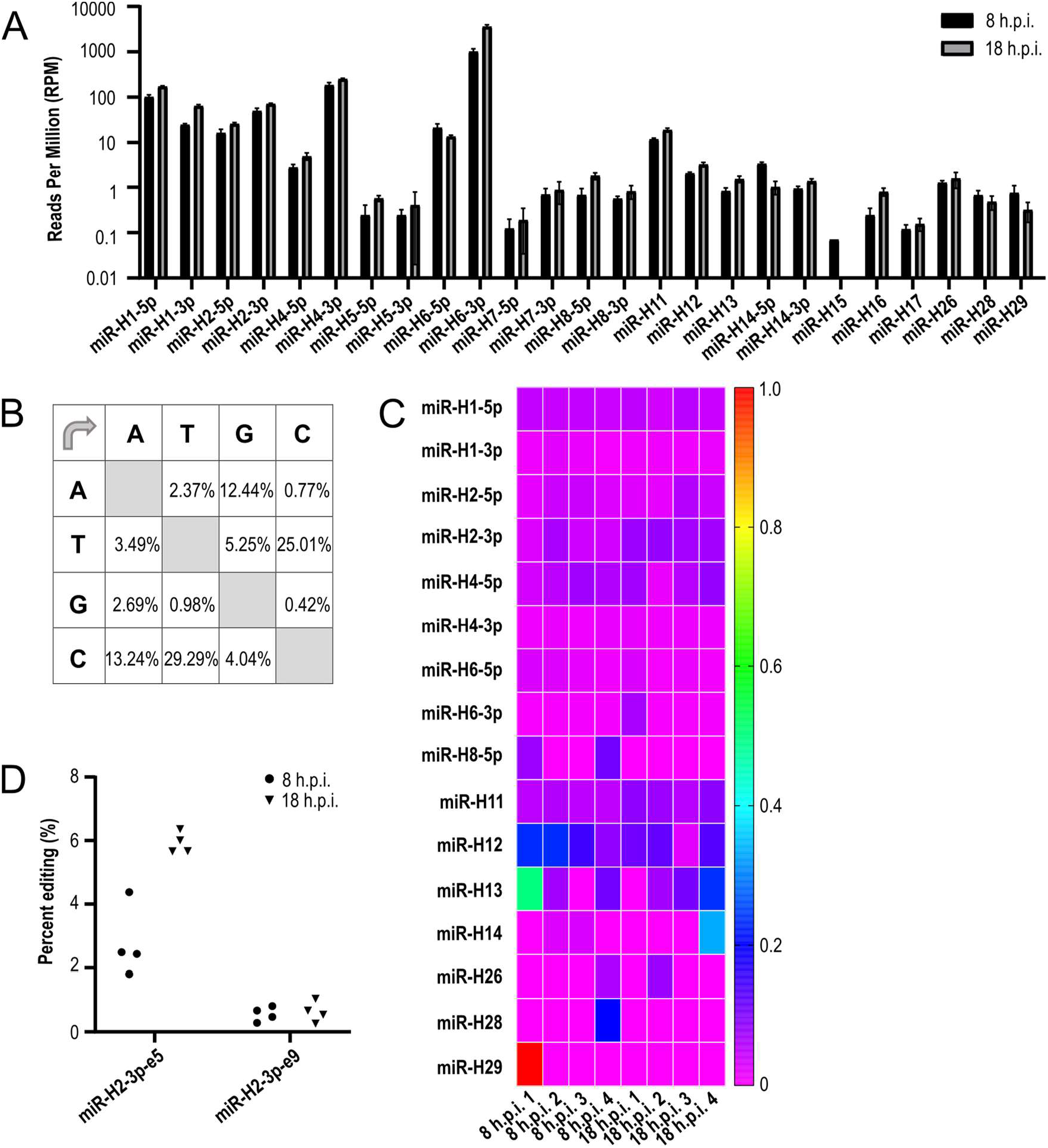
**Low levels of A-to-I editing during productive HSV-1 infection**. A) HFF cells were infected with HSV-1 strain KOS at an MOI of 5 and smRNAs were analyzed by sequencing. The expression of miRNAs is shown as reads per million (RPM) B) Substitution matrix of all detected HSV-1 miRNAs (N=243 714 HSV-1 miRNA reads). C) Each field represents the frequency of A-to-G substitutions for the miRNAs listed below in each sample. D) Percentage of edited miR-H2-3p reads. miR-H2-3p-e5 and miR- H2-3p-e9 indicate sequence editing at the nucleotide position N5 and N9, respectively (number of all miR-H2-3p reads = 273).

Interestingly, similar to latency in humans, we observed an overrepresentation of T-to-C (25.0%) and also a high level of C-to-T (29.3%) nucleotide substitutions, but these were all randomly scattered across all detected miRNAs and had a low frequency (< 1%) (Figure 4B). Although it is possible that some of these represent sequencing errors, it cannot be excluded that C-to-T nucleotide substitutions are the result of random APOBEC3 activity.

### miR-H2 is edited in latently infected mouse trigeminal ganglia

To investigate whether the representative animal model recapitulates the miRNA editing profile in humans, we analyzed miRNAs from acutely and latently infected mouse TGs (3, 14, and 30 days p.i.). As expected, we found an increasing number of miRNAs and reads for each miRNA detected with increasing time post-infection, consistent with the previously determined kinetics of virus replication and establishment of latency in the mouse model after corneal scarification (Figure 5 A). The pattern of miRNAs and nucleotide substitutions within miRNAs resembled the pattern found in human tissue. A-to-G substitutions accounted for the majority of nucleotide substitutions (32%) (Figure 5B), but were not as pronounced as in human tissues. Nevertheless, miR-H2 changes represented the vast majority of these substitutions (72% and 90% at D14 and D30, respectively), and the pattern was similar to that found in humans (ie, miR-H2-e5 10.7% and miR-H2-e9 3.5% at D30) (Figure 5C and D).

**Figure 5.**
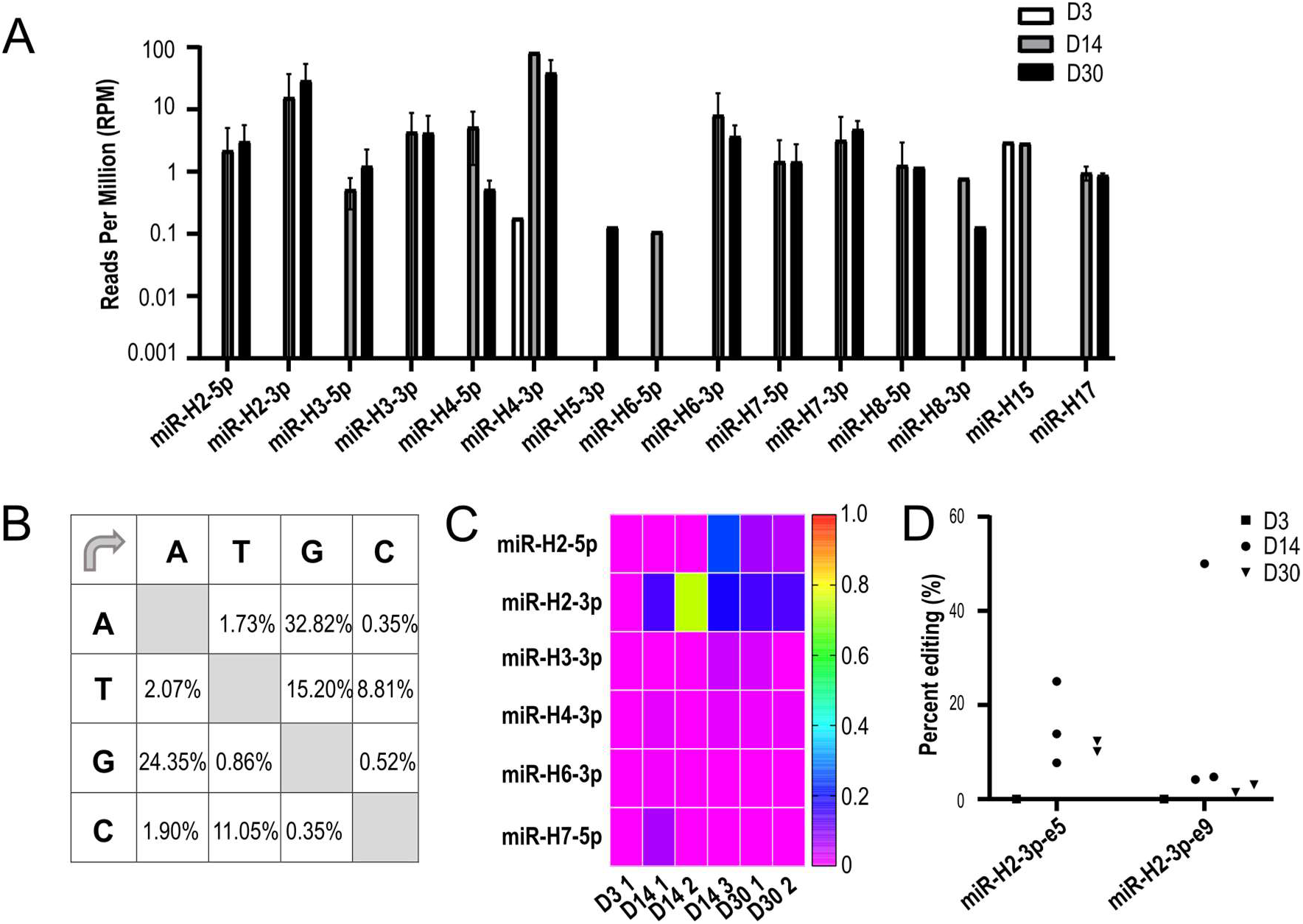
A-to-I hpyerediting during latent HSV-1 infection in mice. A) RNA was extracted from mouse trigeminal ganglia at 3, 14, and 30 days p.i. and small RNA libraries were generated using the NEBNext Multiplex Small RNA Library Pre Set for Illumina and sequenced on the NextSeq 500 (Illumina). The obtained sequencing data sets were processed using sRNAtoolbox. The bars represent the number of reads per million for the miRNAs detected in individual mouse ganglia marked using different colors. B) Substitution matrix of all detected HSV-1 miRNAs (N= 3 858 HSV-1 miRNA reads). C) Each field represents the frequency of A-to-G substitutions for the miRNAs listed below in each sample. D) Percentage of edited miR-H2-3p reads. miR-H2-3p-e5 and miR-H2-3p-e9 indicate sequence editing at the nucleotide position N5 and N9, respectively (Number of reads = 149).

These results indicate that HSV-1 latency in mice, at least in terms of miRNA expression and the ability of mouse proteins ADAR to edit miR-H2, largely recapitulates human latency.

### miR-H2 expressed by HSV-2 does not show evidence of hyperediting

We were also interested in whether hsv2-miR-H2-3p encoded by HSV-2, the sequence and position homolog of hsv1-miR-H2-3p, also shows evidence of editing in productive and latent infection. We use our previously described datasets (9) to reanalyze them and find evidence of hyperdating. Briefly, for analysis of productive infection, HEK-293 cells were infected with the HSV-2 strain 186ΔKpn, a thymidine kinase-negative viral mutant that can establish latency in TGs without lethality to mice, with a MOI of 1, and samples for sequencing were collected 18 h.p.i. For analysis of latency, mice were infected internasally and TGs harvested 30 d.p.i. As expected, we detected a large number of miRNAs during productive infection (Figure 6A) and only a limited number of miRNAs during latency (hsv2- miR-H2 - H7 and -H24). Similar to productive infection of HSV-1, A-to-G substitutions accounted for approximately 12% and 23% of all substitutions in productive and latent infection, respectively (Figure 6B and C). hsv2-miR-H2-3p was detected with 40 reads, but we were unable to detect any read with the A-to-G substitution. We are aware that this is a relatively small number of reads, but we were still surprised that we did not find any read with A-to-G substitutions (Fig 6. D). Interestingly and similar to HSV-1, miRNA arising from a transcript spanning the oriL palindrome, hsv2-miR-H11, shows relatively high frequency of A-to-G substitutions (16,2%) in productive infection (Fig. 6A and B). Overall, the most abundant substitutions in HSV-2 experiment were G-to-A (Fig. 6 B) for the productive infection, G-to-T for the latent sample (Figure 6C). However, we did not observe confident higher representation of specific substitutions for any of miRNAs (i.e. all below 1%), but rather scattered across all detected reads.

**Figure 6.**
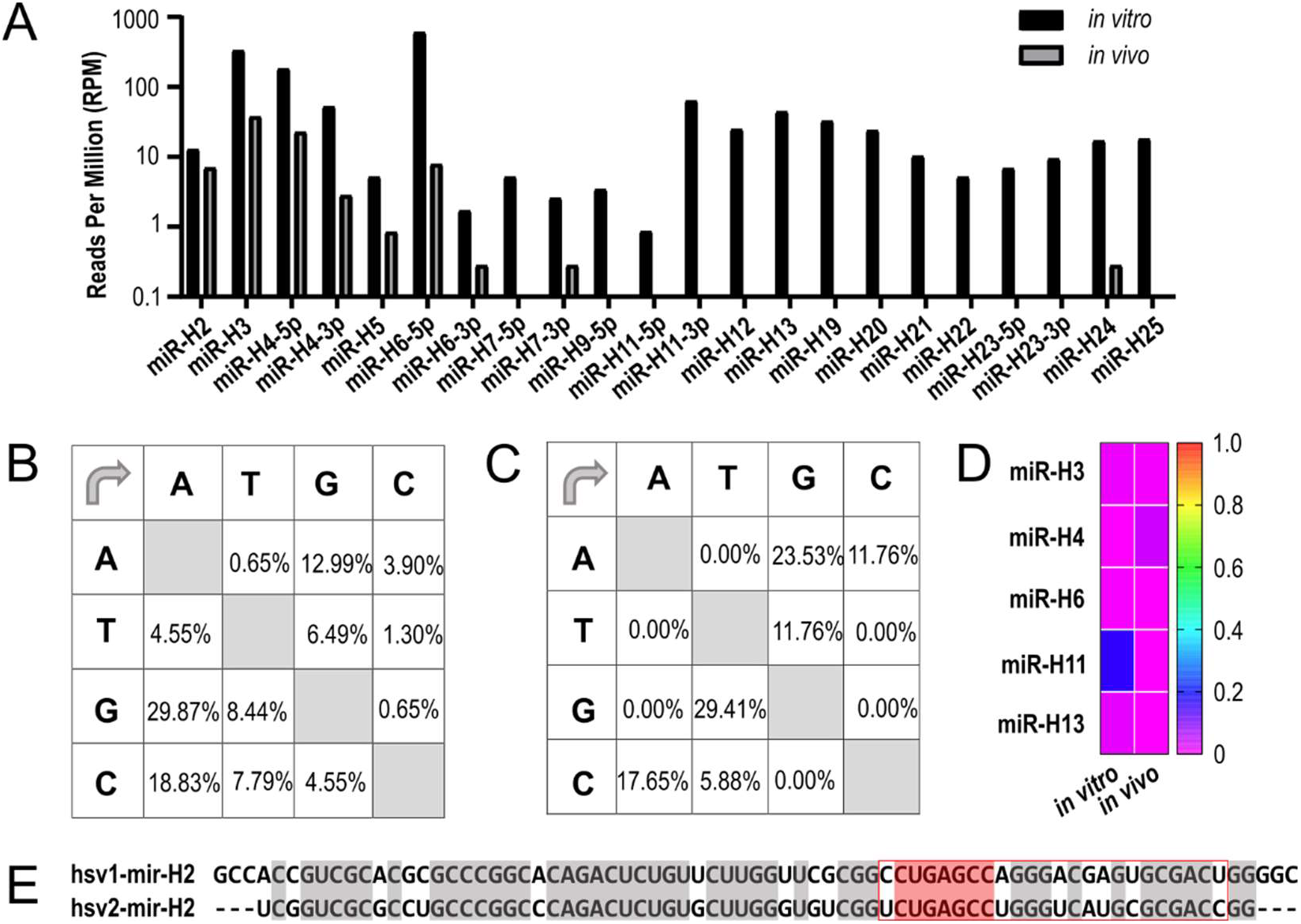
**No evidence for A to I hyperediting in productive or latent HSV-2 infection**. A) HEK293 cells were infected with HSV-2 thymidine kinase-negative mutant of HSV-2 strain 186syn+ (HSV-2) at an MOI of 1 for productive infection, and for the latent infection 7-week-old male CD-1 mice (Harlan) were infected intranasally with 1 × 106 PFU of HSV-2, mTGs were extracted 30 days p.i., and smRNAs were analyzed by sequencing. The expression of miRNAs is shown as reads per million (RPM); B) Substitution matrix of all HSV-1 miRNAs detected in HEK293 samples productively infected with HSV- 2 (N= 1705 HSV-2 miRNA reads). C) Substitution matrix of all HSV-1 miRNAs detected in mTG samples 30 days p.i. latently infected with HSV-2 (N= 284 HSV-2 miRNA reads). D) Each field represents the frequency of A-to-G substitutions for the miRNAs listed below in each sample. E) Sequence comparison between pre-mir-H2 of HSV-1 strain 17 and HSV-2 strain HG52. The miR-H2-3p is shown in the red frame; and the seed sequence is shown in red.

### Editing of miR-H2 extends its targeting potential

We show evidence of miRNA editing in latently infected human ganglia and in latently infected mice. Moreover, editing can also be observed during productive infection in culture, the system in which miRNA play a minor role in efficient viral replication. Overall, it is expected that it will be very difficult to determine the biological significance of these events using current experimental models. miR-H2 deletion mutants, for example, show little phenotype in mice (25). Nonetheless, we wanted to investigate the potential of edited miRNAs to regulate additional viral or host targets.

First, we asked whether the edited form is efficiently associated with the RISC complex, which should indicate their inhibitory potential. To this end, we infected HFF cells and extracted 12 h.p.i. RNAs from whole cell lysates or immunoprecipitated Ago2, the major component of the RNA-inducing silencing complex (RISC). We postulated that a consistent ratio of canonical to edited miRNA in Ago2 pulldown and total miRNA would indicate efficient loading of edited miRNA into RISC (49). Indeed, the percentage of edited miRNA (miR-H2-3p-e5) in the total miRNA samples was 8.09%, while it was 10.23% in the Ago2 pulldown, indicating an efficient association with RISC.

Next, we predicted the host and viral targets of these miRNAs. Not surprisingly, the targets of miR-H2 and miR-H2-e5 overlapped only slightly using two common prediction algorithms, TargetScan and miRDB (Figure 7 A ad Suppl. Table 4). These results suggest that editing significantly increases the potential of miR-H2 to regulate host genes. Of note, miR- H2-e9 was not included in the analysis because the host target prediction algorithms are largely based on the seed sequence, and miR-H2-e9 has the same sequence as miR-H2. However, this does not exclude a possibility that miR-H2-e9 regulated different transcripts.

**Figure 7.**
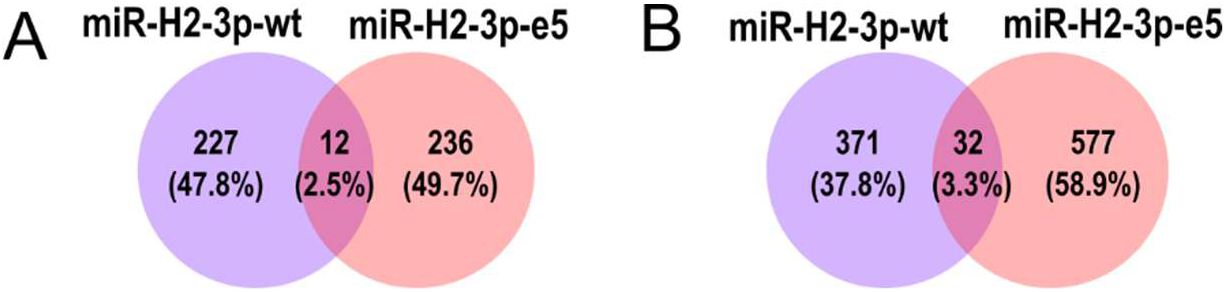
Targeting potential of miR-H2 is expanded by editing of the seed sequence. Predicted human targets of canonical (purple) miR-H2-3p and edited miR-H2-3p (orange) using: A) TargetScan ((66)); and B) miRDB (64).

Regarding potential viral targets, miR-H2 is fully complementary to the ICP0 transcript and likely represents the main target of miR-H2 (8). We used the Bowtie alignment tool (50) to predict potential viral targets of miR-H2 and its edited forms. We identified 17 potential binding sites for miR-H2-3p, including the previously identified ICP0. Surprisingly, one nucleotide substitution within the seed region of miR-H2 (i.e. miR-H2-e5) results in a dramatic increase in predicted targets (i.e., 79 loci), which include ICP0, ICP4, and a large number of other productive infection gene transcripts (Suppl. Table 2). Although functional characterization of miR-H2-e5 is experimentally challenging and beyond the scope of this manuscript, we carefully analyzed the potential binding sites for miR-H2 and miR-H2-e5 within ICP0 and ICP4 transcripts using RNAhybrid. Similar to Bowtie prediction, editing miR- H2 increased the number of potential target sites in both transcripts (Suppl. Table 3). Indeed, the canonical miR-H2 has only one fully matched seed-target site in the ICP0 and ICP4 transcripts, whereas editing at position N5 results in 4 and 7 potential seed matches in the ICP0 and ICP4 transcripts, respectively.

To preliminary test whether miR-H2-3p edited at positions 5 and/or 9 had the potential to regulate the expression of ICP0 or ICP4, we performed co-transfection experiments. Briefly, we co-transfected HEK293 cells with a wild-type ICP0 or ICP4 expression plasmid together with a pEGFP plasmid (an indicator of non-selective targeting) and microRNA mimics including: canonical miR-H2-3p, miR-H2-3p with scrambled seed sequence (miR-H2-3p-mut), negative control mimic (scrambled sequence), edited miR-H2-3p at position 5 and/or 9, or both (miR-H2-3p-e5, miR-H2-3p-e9, miR-H2-3p-e5-e9 (Fig. 8). As shown in Figure 8, canonical miR-H2-3p significantly decreased ICP0 levels but not ICP4 levels. On the other hand, miR-H2-e5, but not miR-H2-e9 and miR-H2-e5-e9, decreased the levels of ICP0 and ICP4 (Figure 8 B and C). The levels of these proteins were not affected by miR-H2-3p-mut and negative control. These initial results, albeit in an overexpression system, suggest that miR-H2 may play a broader role in repressing early genes during latency.

**Figure 8.**
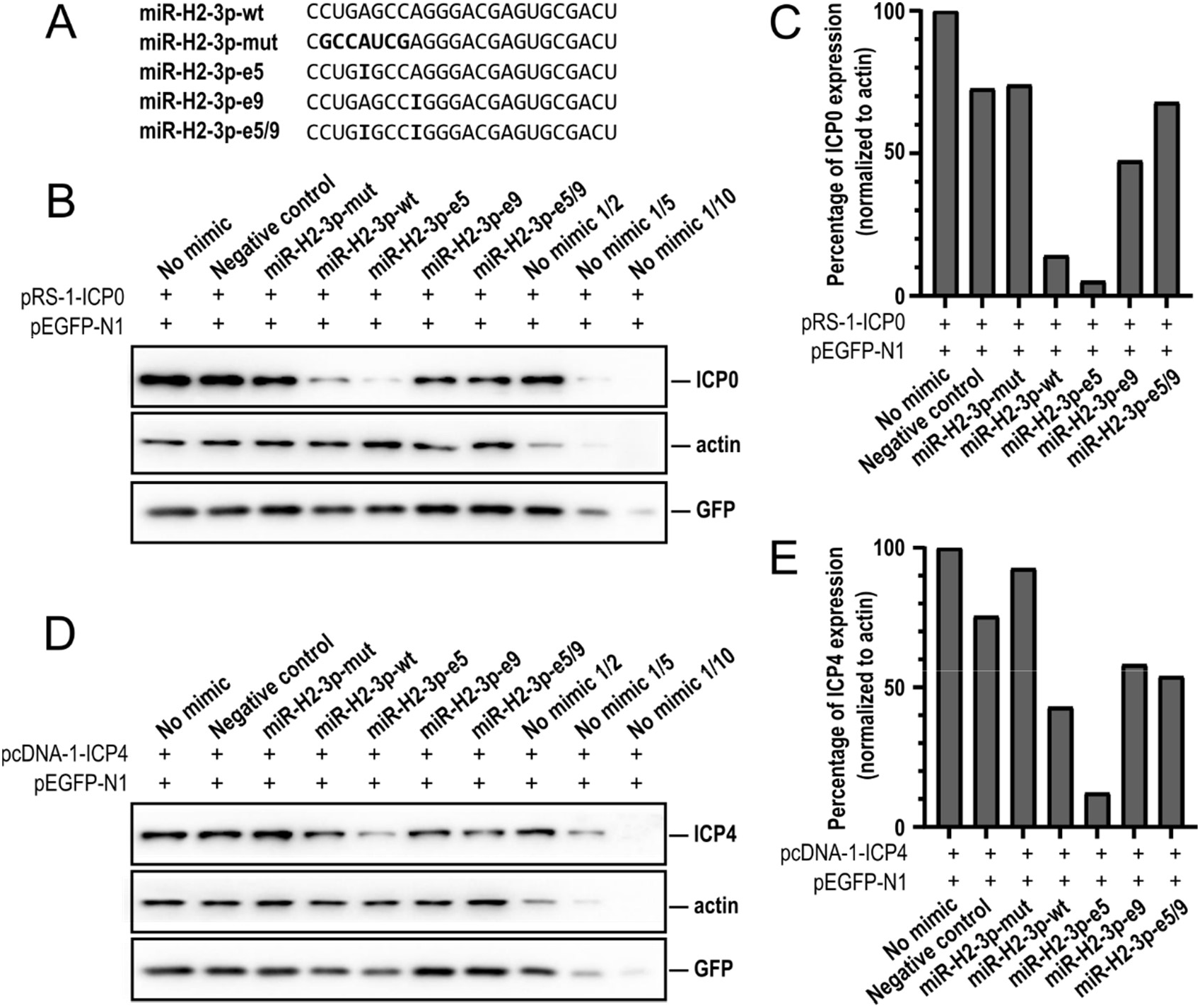
miR-H2-3p-e5 can downregulate ICP0 and ICP4. A) Sequences of miRNA mimics. Single stranded mimics with modified 2’Ome corresponding the canonical or edited sequence were used: canonical miR-H2-3p (miR-H2-3p-wt), a mutated seed sequence of the miR-H2-3p marked bold (miR- H2-3p-mut), edited miR-H2-3p on positions 5 and/or 9 or both (miR-H2-3p-e5, miR-H2-3p-e9, miR-H2- 3p-e5-e9), inosines are marked bold. B) Proteins extracted from HEK293 cells transfected with plasmid expressing GFP (pEGFP-N1) and ICP0 (pRS-1-ICP0), and co-transfected with miRNA mimics indicated above the panel, were analyzed by Western blot. C) Quantification of ICP0 protein levels from B) using ImageJ, and normalized to actin; D) HEK293 cells were transfected plasmids expressing GFP (pEGFP- N1) and ICP4 (pcDNA-1-ICP4), and co-transfected with miRNA mimics indicated above the panel. E) Quantification of ICP4 protein levels from D) using ImageJ, and normalized to actin.

## Discussion

Our data show that miR-H2 encoded by HSV-1 is post-transcriptionally edited in latently infected human neurons, leading to an expansion of potential host and viral targets that may play a role in regulating latency. The observed A-to-G substitutions are the signature of the ADAR proteins that catalyze the deamination of pri-/pre-miRNAs. Because of relatively small number of latently infected neurons, HSV-1 miRNAs are present at very low levels in human tissues, so our analysis is largely based on next-generation sequencing, which can provide sufficient depth and resolution.

### The miRNA pattern of HSV-1 latency

Our analysis of ten latently infected human ganglia reveals remarkably similar patterns of HSV-1-encoded miRNAs. These include miR-H2 and miR-H8, which likely determine the HSV-1 miRNA pattern. We cannot rule out the possibility that with increasing depth (higher number of reads) other miRNAs are not found, but would certainly be underrepresented. Similar to humans, latently infected ganglia from mice have the same miRNA expression pattern. It is important to note that standard latent infection models in mice are based on 30 days infection, which is far less that latency studied in humans. Also, latency in mice is established after massive productive infection within TG, which is not the case in humans. Therefore, there is increased potential for residues of productive infection transcripts to exist at 30-day post-infection mice compared to the human samples, including appearance of some miRNA. The exact function of these miRNAs is still quite enigmatic, mainly because studies on HSV-1 miRNAs in during the latent phase of infection has multiple challenges and drawbacks. An important advance in understanding the role of HSV-1 miRNAs has been made using viral mutants lacking single miRNAs, but the observed phenotypes are generally mild (21, 25–27, 51, 52). This is to be expected because miRNAs likely regulate the robustness of the system (i.e., sensitive on/off switches) rather than enabling latency or reactivation. Interestingly, in HSV-1 miRNAs present in latency have both arms of the miRNA duplex, suggesting that viruses use the miRNA machinery particularly efficiently.

### Editing of miR-H2

Our study provides several lines of evidence that miR-H2 is hyperedited during latency, further increasing its targeting potential. First, the sequencing data were analyzed under stringent cutoff values, resulting in highly reliable sequence information. Second, sequencing of DNA in all samples analyzed revealed no evidence of heterogeneity within the pre-miR-H2 locus, suggesting that miR-H2 heterogeneity is due to post-transcriptional modifications of its transcripts. Third, the pattern of miR-H2 expression (ratio of edited to unedited forms) was remarkably consistent in all samples. It is highly unlikely that all patients carry similar genomic variants. Fourth, evidence from sequencing of numerous viral genomes suggests that the miR-H2 locus is highly conserved. Fifth, we found the same editing pattern (i.e. miR-H2 positions N5 and N9) in productively infected cells in culture, albeit at lower frequency, and in mouse model, suggesting a specific editing process. We believe that the lower frequency of miR-H2 editing in productive infection is due to the ADAR proteins being saturated with the overwhelming viral transcript. This is in contrast to the latency phase, where fewer transcripts are present and can be efficiently edited.

Interestingly, we found no evidence of site-specific hyperediting for the encoded HSV-2 homolog; however, in the context of limitations this study, we cannot rule out the possibility that hsv2-miR-H2 is not edited, especially as mice were infected with a TK-null virus, which would limit viral replication in the ganglia prior to latent establishment. In the HSV-2 dataset, we found relatively small number of reads representing miR-H2, and unfortunately, we were unable to obtain human tissues latently infected with HSV-2.

How the ADAR1 and ADAR2 editors select their targets is not clear, but the length and structure of the dsRNA (bulges, loops, and mismatches) play an important role. The precursors of the pre-miRNAs of HSV-1 and HSV-2 are very similar (9), and the adenosine targeted by editing at position N5 is conserved, and we found no clear evidence or motifs for differential specificity of editing. For the precursor of HSV-1 miR-H2 (pre-miR-H2), the optimal secondary structure was predicted to have a slightly higher minimum free energy (- 37.50 kcal/mol), higher than all other commonly expressed miRNAs and higher than HSV-2 miR-H2 (-40.40 kcal/mol). Editing at the N5 position slightly lowers the folding energy.

Nevertheless, detailed studies on host miRNAs and viral miRNAs are needed to establish such a correlation. In addition to the A-to-G substitutions, we observed relatively many other substitutions in our analysis. Some of these could be due to technical sequencing errors (47), while others could be active processes. The APOBEC (apolipoprotein B mRNA editing catalytic polypeptide-like) family of proteins bind to both RNA and single-stranded (ss) DNA and may be responsible for some of the C-to-T substitutions (53), but investigating this is beyond the scope of this study.

### Possible roles of editing

The main question is whether editing of miR-H2 has biologically relevant functions. Our results indicate that the edited miRNA is loaded into the RISC complex as efficiently as the canonical sequence, suggesting functional properties.

Furthermore, we show that editing miR-H2, similar to other edited miRNAs, increases its targeting potential. However, viruses lacking miR-H2 expression, which includes both canonical and edited miRNA, exhibit a mild phenotype in latency models (25, 27, 51), suggesting that editing may at best contribute to the observed phenotype. Current latency models rely on relatively harsh experimental conditions, including viral reactivation in explanted tissues, which may not provide the resolution necessary to study miRNA-regulated processes. However, recently developed models in primary neurons of latency establishment and reactivation, where reactivation is induced on intact neurons, may in the future prove useful for investigating the function of ADAR in HSV neuronal infection (54). One potential caveat however is that studying the biological role of editing is challenging because it is challenging to generate ADAR1 KO in most cells. In addition to the evidence that editing affects targeting properties, it is also possible that editing of the miR-H2 precursor affects its biogenesis, as shown for EBV- and KSHV-encoded and edited miRNAs (39, 41, 42). ADAR1 was shown to directly interact with DICER and affect miRNA processing, maturation, RISC loading, and silencing of target RNAs, all of which remain to be investigated. To our surprise, we did not find evidence of HSV-2 miR-H2 editing, nor in infected cells in culture nor in latently infected trigeminal ganglia. At this stage, because of the relatively small number of reads, we cannot rule out the possibility that hsv2-miR-H2 is edited. We can speculate that editing may be the molecular mechanism that determines some biological differences between HSV-1 and HSV-2. Therefore, it is critical to sequence relevant human samples, such as the dorsal root ganglia, to determine the status of HSV-2 miRNAs during latency. In addition, our study suggests that other noncoding RNAs, i.e. LAT intron, can be edited during latency, which may explain some of the peculiar properties of this molecule. For example, editing the LAT intron could prevent this abundantly expressed molecule from being recognized as non-self, which would lead to apoptosis (55). Editing could also affect its expression and stability or be essential for its interactome (56), and yet there is still much to learn about the importance of post-transcriptional editing in herpesvirus infection.

## Materials and methods

The Ethics Committee of the Faculty of Medicine in Rijeka, University of Rijeka approved the use of autopsy samples for the present study. Trigeminal ganglia (TG) were removed from 10 subjects within 24 hours after death (Supplemental Table 1). At the time of death, patients had no symptoms of HSV-1 infection. TG specimens were immediately frozen and stored at - 80°C until further processing.

### Cells, viruses, infection, and plasmids

Human Embryonic Kidney (HEK293, CRL-1573) and Human Foreskin Fibroblast (HFF, generous gift of S. Jonjić, Faculty of Medicine, University of Rijeka) were grown in Dulbecco’s Modified Eagle Medium (DMEM) (PAN-biotech) supplemented with 10% fetal bovine serum (FBS; PAN-biotech), Penicillin / Streptomycin 100 μg / μl, 2 mM L-glutamine (Capricorn), and 1mM Sodium Pyruvate (Capricorn), under standard conditions in humidified incubator at 37°C and in the presence of 5% CO2. Wild-type HSV-1 strain KOS (kindly provided by Professor Donald M. Coen and David Knipe, Harvard Medical School, Boston, USA) was prepared in Vero cells and stored at -80°C as previously described (57). For productive infection, cells were seeded on the dish or plate the day before the experiment. After 24 hours, cells were infected with specified viruses from a viral stock, with indicated MOI (multiplicity of infection) or mock infected (uninfected). The medium was in all samples replaced with fresh growth medium 1 hour after infection. Samples were collected at specified hours post infection (h.p.i.). For latent infection, 6-week-old female CD-1 mice (Charles River) were anesthetized by intraperitoneal injection of ketamine hydrochloride (80 mg/kg of body weight) and xylazine hydrochloride (10 mg/kg) and inoculated with 1.5 × 10^6 PFU/eye of KOS strain HSV-1 (in a 5- μL volume) onto scarified corneas, as described previously (58). Mice were housed in accordance with institutional and National Institutes of Health guidelines on the care and use of animals in research, and all procedures were approved by the Institutional Animal Care and Use Committee of the University of Virginia. Trigeminal ganglia were harvested 3, 14 and 30 days after infection and immediately snap frozen in liquid nitrogen. For RNA extraction, lysis was carried out by addition of RNA lysis buffer (Zymo) and homogenization for 60 s using the BeadBug^TM^ microtube homogenizer. RNA was isolated from the homogenized mixture using the *Quick*-RNA miniprep kit (Zymo).

The ICP0-WT plasmid (pRS-1) was a generous gift of Rozanne Sandri-Goldin (University of California). This plasmid includes the entire *ICP0* gene, with its promoter, cloned into pUC-18 (31, 59). pcDNA-1-ICP4 contains the entire *ICP4* gene (60). pEGFP-N1 was obtained from Clontech-Takara.

### Nucleic acid extraction

Trigeminal ganglia were thawed and small sections homogenized and dissolved in TRIreagent (Ambion). DNA and RNA were extracted from the same sample according to the manufacturer’s instructions. Briefly, chloroform was added in TRIreagent to separate the RNA in the aqueous phase and precipitated with Isopropanol. After washing with 70% ethanol, precipitate was resuspended in nuclease free water and treated with DNase for 30 min. The remaining sample from which the aqueous phase was removed was used for DNA precipitation with absolute ethanol. HEK293T and HFF samples were homogenized in TRIReagent solution (Ambion) on ice and total RNA was extracted according to the manufacturer’s protocol. DNA and RNA concentration and quality were measured using UV/VIS spectrophotometer (BioDrop μLITE, UK).

### Massive parallel sequencing

Small RNA sequencing libraries were prepared using NEBNext Multiplex Small RNA Library Prep Set for Illumina using standard protocol according to the manufacturer’s instructions. Libraries were size selected by using AMPure XP beads to selectively bind DNA fragments 100 bp and larger to paramagnetic beads. Samples were sequenced on NextSeq 550 NGS platform (Illumina) using single end sequencing mode with total of 36 cycles. RNAs extracted from immunoprecipitated complexes using Ago2 and GFP antibody were used to generate small RNA sequencing libraries using TruSeq Small RNA Kit (Illumina), according to the manufacturer’s instructions. cDNA libraries were size selected using 10% PAGE (BioRad), and species from 15 to 30 nt were isolated. cDNA libraries were then sequenced by single read sequencing (50 nt) in Illumina HiSeq 2500 sequencer.

### Data analysis

For quality control check on raw sequences, we used FastQC. The sequencing data was analyzed using sRNAbench (Computational Epigenomics Lab, Evolutionary Genomics and Bioinformatics Group. Dept. of Genetics, Inst. of Biotechnology, University of Granada, Spain) (45, 61, 62). Briefly, 3’ flanking sequencing adapter sequence was removed from all the reads, and reads were aligned against the human reference genome (Genome Reference Consortium Human Build 38 patch release 13) and HSV-1 strain 17 (NC_001806.2) or HSV-1 KOS strain (JQ673480.1). Reads smaller than 18 were excluded, as were the reads with a sequencing quality score lower than 30, to obtain high-quality reads. To assign the reads to known miRNAs, reads were aligned against known miRNAs from the miRBase sequence database (release 22), human and HSV-1, while published HSV-1 miRNAs not represented within the miRBase, hsv1-miR-H28, and hsv1-miR-H29, were added to the database file based on the published sequence. We allowed for 2 nucleotide mismatches in the first 19 nucleotides mapped but also accepted all sequences that started at most 3 nucleotides upstream and ended at most 5 nucleotides downstream of the reference sequence.

For the prediction of the host targets, prediction tools such as TargetScanHuman 5.2 Custom (63)and miRDB (64) were used to search for the presence of sites that match the seed region of miRNA, nucleotides from 2 to 8, and for the viral targets, we also searched for the presence of sites complementary to the seed region of miRNA using Bowtie alignment tool (50). In addition, we used RNAhybrid (65) applying standard settings to visualize 10 miRNA-target bindings in the ICP0 and ICP4 region.

### Immunohistochemistry

Immunohistochemical staining of ADAR proteins was conducted using DAKO EnVision+System, Peroxidase (DAB) kit (DAKO Cytomation, Santa Clara, CA, USA) on 4 µm thick serial sections of paraffin-embedded TG tissues. Briefly, after deparaffinization and rehydration, tissues underwent heat-mediated epitope retrieval by microwave heating in a 10-mM citrate buffer, pH=6.0. Having subsequently been treated with blocking solution, slides were incubated with rabbit monoclonal α-ADAR1 IgG (Cell Signaling Technology; diluted 1:100 in 1% BSA in PBS), mouse monoclonal α-ADAR1 IgG (Santa Cruz; diluted 1:100 in 1% BSA in PBS) and mouse monoclonal α-ADAR2 IgG (Santa Cruz; diluted 1:100 in 1% BSA in PBS) over 12 hours in a humid chamber at 4°C. As a secondary antibody, a peroxidase-labeled polymer linked to goat α-rabbit and α-mouse immunoglobulins was applied for 30 min at room temperature. Immunoreactions were visualized by 3,3′-Diaminobenzidine (DAB) and slides were counterstained with hematoxylin. After being dehydrated, slides were mounted with Entellan (Sigma-Aldrich, Hamburg, Germany) and analyzed by an Olympus BX51 microscope equipped with a DP50 camera and Cell^F software (Olympus, Japan). The specificity of antibody binding was verified performing negative controls by substitution of ADARs antibodies with isotype-matched control antibodies applied in the same conditions. Negative control slides showed no immunohistochemical signals.

### Transfection

Single stranded RNAs with modified 2’Ome designed to mimic canonical miR-H2-3p (miR-H2- 3p-wt - 5’- CCUGAGCCAGGGACGAGUGCGACU-3’), mutated seed sequence of the miR-H2-3p (miR-H2-3p-mut - 5’- CGCCAUCGAGGGACGAGUGCGACU-3’), edited miR-H2-3p on positions 5 and/or 9 or both (miR-H2-3p-e5 – 5’- CCUGIGCCAGGGACGAGUGCGACU-3’, miR-H2-3p-e9 – 5’- CCUGAGCCIGGGACGAGUGCGACU-3’, miR-H2-3p-e5-e9 – 5’- CCUGIGCCIGGGACGAGUGCGACU-3’), and negative control mimic were ordered from GenePharma. Plasmid (pRS-1-ICP0 / pcDNA-1-ICP4 / pEGFP-N1) and miRNA mimic co-transfections were performed in HEK293T cells using Lipofectamine 3000 according to the manufacturer’s instructions without the addition of the P3000 reagent. Briefly, cells were seeded in 24-well plate to be 70% confluent for the transfection. Cells were co-transfected with 70 ng per well of ICP0 expression plasmid or 50 ng per well of the ICP4 expression plasmid, 30 ng per well of the pEGFP-N1 expression plasmid, and with 30 nM of the mimic or negative control, using one microliter of Lipofectamine 3000 per well. After 24 hours, samples were extracted for Western blot analysis.

### Western blot

To extract the proteins, cells were collected at indicated timepoints and lysed in RIPA buffer (150 mM NaCl, 1% NP-40, 0.5% Na deoxycholate, 0.1% SDS, 50 mM Tris (pH 8.0), with protease inhibitors (cOmplete, Roche, Basel, Switzerland)), mixed with 2x Laemmli buffer with β- mercaptoethanol (Santa Cruz Biotech, Dallas, TX, USA), and denatured for 6 min at 95°C. Proteins were separated in 10% SDS–PAGE gels and transferred onto a nitrocellulose membrane (Santa Cruz Biotech, Dallas, TX, USA). Membranes were blocked in 5% w/v nonfat dry milk in 1x Tris-buffered saline (TBS) for 30 min at room temperature followed by the incubation at 4°C with gentle rotation overnight with the primary antibodies with specified dilutions: α-actin (MilliporeSigma, Burlington, MA, USA)—1:10000, α-ICP0 (Abcam, Cambridge, UK)—1:2000, α-ICP4 (Abcam, Cambridge, UK)—1:2000, α-ADAR1 (Santa Cruz Biotech, Dallas, TX, USA) – 1:1000, α-ADAR1p150 (Cell Signaling Technology, Inc., Danvers, MA, USA) – 1:1000. Blots were washed for 30 min with TBS-0.05% Tween 20 (TBS-T) and primary antibodies were detected using horseradish peroxidase-conjugated goat anti-mouse secondary antibody diluted 1:2000 (Cell Signaling Technology, Inc., Danvers, MA, USA) and incubated at room temperature for 1 h. Blots were again washed for 30 min and visualized using the Amersham ECL reagent or SuperSignal West Femto Maximum Sensitivity Substrate (Thermo Fisher Scientific, Waltham, MA, USA) and ChemiDoc MP (Bio-Rad Laboratories, Hercules, CA, USA).

### Immunoprecipitation

HFF cells were seeded in 10 cm dishes in triplicates for the infection with HSV-1 KOS strain and extracted at 12 h.p.i. From each sample, a fraction (1/10) was taken for the total population of miRNAs before the immunoprecipitation protocol, and the rest of the sample was used for Ago and GFP immunoprecipitation (control). Immunoprecipitation of RNA bound to Argonaute (Ago) proteins was performed according to the Dynabeads Protein G protocol (Thermo Fischer Sc.). Briefly, Dynabeads were incubated with the 10 μg of α-Ago antibody (Millipore), or α-GFP (Millipore) that acted as a negative control, diluted in PBS with 0,1% Tween20 overnight at 4°C. After incubation, the supernatant was removed using a magnet, and beads were washed with PBS with 0,1% Tween20. HFF samples were collected 12 hpi in triplicates using NP-40 lysis buffer, and supernatant from cell lysis was added to the beads-Antibody complex and incubated overnight at 4°C. After incubation, beads were washed and resuspended in TriReagent solution, and total RNA was extracted according to the manufacturer’s instructions.

## Author Contributions

Conceptualization I.J.; methodology I.J., M.C. H.J., O.V. and M.H.; validation I.J., M.H., A.C., A.G, O.V. and D.C.; investigation A.Z., A.P., M.C., F.R., H.J., C.G., A.W., S.D. and A.C.; resources, I.J., M.H., A.C., A.G., D.C. and O.V.; writing—original draft preparation, I.J.; writing—review and editing all authors.; supervision, I.J., A.C. D.C: M.H., A.G. and O.V.; project administration, I.J.; funding acquisition, I.J. and M.H.. All authors have read and agreed to the published version of the manuscript.

## Acknowledgements

The study was supported with Croatian Science Foundation Grant IP-2020-02-2287 and DOK- 2012-02.9152, and University of Rijeka support grant prirod-sp-23-502930 to IJ; AIRC IG grant (n. 13202) to A.G.; and National Institutes of Health grants NS105630 to A.R.C., T32GM008136 to S.A.D. and T32AI007046 to A.L.W.

